# *Mycobacterium tuberculosis* infection boosts B cell responses to unrelated pathogens

**DOI:** 10.1101/680058

**Authors:** Simon G. Kimuda, Irene Andia-Biraro, Ismail Sebina, Moses Egesa, Angela Nalwoga, Steven G. Smith, Bernard S. Bagaya, Jonathan Levin, Alison M. Elliott, John G. Raynes, Stephen Cose

**Author notes:** Division of Epidemiology and Biostatistics, School of Public Health, University of the Witwatersrand, Johannesburg, South Africa (Jonathan Levin).

## Abstract

Antigens from *Mycobacterium tuberculosis* (*M.tb)*, have been shown to stimulate human B cell responses to unrelated recall antigens *in vitro*. However, it is not known whether natural *M.tb* infection or whether vaccination with the related species, *Mycobacterium bovis* BCG, has a similar effect. This study investigated the effects of *M.tb* infection and BCG vaccination on B cell responses to heterologous pathogen recall antigens. Antibodies against several bacterial and viral pathogens were quantified by ELISA in 68 uninfected controls, 62 individuals with latent TB infection (LTBI) and 107 active pulmonary TB (APTB) cases, and 24 recently BCG-vaccinated adolescents and naive controls. Antibody avidity was investigated using surface plasmon resonance and B cell ELISPOT assays were used to measure plasmablast and memory B cell responses (MBC) in APTB cases and healthy donor controls. APTB was associated with higher levels of antibodies to tetanus toxoid (TT), diphtheria toxoid, respiratory syncytial virus, measles virus and Kaposi’s sarcoma herpesvirus, compared to uninfected controls. Vaccination with BCG did not alter levels of antibodies against heterologous pathogens. TT-specific antibody avidity was increased in APTB and the ratio of TT-specific plasmablasts to MBCs in the APTB cases was 7:1. *M.tb* infection boosts serological memory to heterologous pathogens in human subjects and this process may be driven by polyclonal activation of memory B cells.

**Significance:** *Mycobacterium tuberculosis (M.tb)* has potent immunostimulatory properties and has been used in adjuvant preparations to improve vaccine responses in animals. This study shows that natural *M.tb* infection in humans is associated with increased antibody and B cell recall responses to heterologous pathogens. This data suggests a potential role for *M.tb* antigens in immunotherapies designed to maintain antibody immunity to diverse infections.

## 1.0 Introduction

The *Mycobacterium tuberculosis* complex (MTBC) is made up of several mycobacterial species that cause tuberculosis (TB), a disease that affects millions of people worldwide. There were 10 million cases of TB disease in 2017, 1.6 million deaths, and a quarter of the world’s population is estimated to be infected (1). Despite the seriousness of the disease caused by MTBC, some of these pathogens have use as immunotherapies due to their potent immunostimulatory properties. The most important member of the MTBC, *Mycobacterium tuberculosis* (*M.tb*), is well known for its adjuvant properties. It is a constituent of Freund’s complete adjuvant and is thought to improve vaccine responses through its stimulatory effect on antigen presenting cells such as dendritic cells and macrophages (2). *Mycobacterium bovis* Bacille Calmette Guerin (BCG) has been used to treat bladder cancer for over 30 years (3). The exact mechanism of action has not been fully elucidated, but the common hypothesis is that BCG draws innate immune cells into the bladder and primes them to attack cancer cells (4).

Increasing evidence has been put forward supporting a potential role for BCG in the protection of infants from diseases caused by heterologous pathogens. Early studies in Guinea Bissau showed a decrease in deaths from infectious diseases among low birth weight infants who were given BCG vaccine at birth (5,6). These effects are thought to be as a result of improved function of innate immune cells brought about by cellular epigenetic modifications induced by BCG (9). There may also be non-specific effects on the adaptive immune system. Ota *et al*. (7) described higher hepatitis B virus-specific antibody responses in infants who were given BCG in addition to hepatitis B vaccination at birth, compared to those who were only vaccinated with hepatitis B. A more recent study by Ritz and colleagues (8) reported that the level of antibodies elicited by pneumococcal vaccination of infants at two, four and six months of age were higher in those infants who were given BCG at birth compared to those who were not. BCG may therefore improve immune responses to vaccines given at the same time as BCG or those given following BCG vaccination, in a non-specific manner.

The studies highlighted above describe the effect of mycobacteria on responses to concurrently administered vaccine antigens or those given after vaccination with mycobacterial preparations. However, studies in the past have shown that purified protein derivative (PPD) from *M.tb* can stimulate secretion of antibodies against measles, rubella and herpes simplex viruses from human peripheral blood mononuclear cells (PBMCs) *in vitro* (10). This finding suggests that *M.tb* may be able to enhance antibody/B-cell memory responses generated from previous exposure to unrelated pathogen-derived antigens. The exact immunological mechanism underlying this observation is not yet known; however, it is possible that *Mycobacterium* antigens could non-specifically activate pathogen-specific memory B cells (MBCs), resulting in the expansion of antibody-secreting cells and a subsequent rise in antibody levels. Human MBCs are prone to activation by polyclonal stimulation; studies by Bernasconi and colleagues (11) have shown that stimulation of these cells by bacterial CpG (cytosine-phosphate-guanine) DNA or by T cell cytokines can lead to their proliferation and expansion into antibody-secreting cells. However, this hypothesis has not been investigated in regard to *Mycobacterium tuberculosis* exposure. Furthermore, it is not known whether natural *M.tb* infection has the same non-specific stimulatory effect on antibody responses to recall antigens from heterologous pathogens. We also do not know whether recent vaccination with the related species, *M. bovis* BCG has a similar effect on serological recall responses to unrelated antigens.

This study characterised antibody responses to heterologous pathogen recall antigens in uninfected controls, individuals with a latent TB infection (LTBI) or active pulmonary TB cases (APTB) cases participating in a TB household contact study in Uganda, as well as adolescent recipients of the BCG vaccine and their age-matched naïve controls from the United Kingdom. Additionally, polyclonal activation of MBCs was explored as a possible mechanism by studying frequencies of tetanus toxoid (TT)-specific plasmablasts and MBCs in the APTB cases and healthy donors.

## 2.0 Results

### 2.1 Demographics

The characteristics of the Ugandan TB household contact study participants have been described previously (12) and in Supplementary Table 1. In summary, APTB cases and individuals with LTBI were older than the uninfected individuals and there was a larger proportion of males in the APTB group in comparison to the other two groups. There were also more HIV positive individuals and a larger proportion of people of low socioeconomic class among the APTB group.

The adolescents studied in the UK were all aged between 12 and 13 years, had no previous BCG scar and were tuberculin skin test negative (13).

### 2.2 *M.tb* infection state is associated with increased antibody responses to unrelated pathogens

Antibody responses were compared across groups of uninfected individuals, individuals with LTBI and APTB cases from Uganda to determine the impact of *M.tb* infection state on antibody responses to heterologous pathogens. Since we were interested in recall responses, we studied antibodies against antigens contained in childhood vaccines, such as tetanus toxoid (TT), diphtheria toxoid (DT) and measles virus (MV). We also measured antibodies specific for infections that our participants may have had a high chance of encountering in childhood in this setting. These included respiratory syncytial virus (RSV) (14), adenovirus (15), Kaposi’s sarcoma herpes virus (KSHV) (16), cytomegalovirus (CMV) (17) and Epstein Barr virus (EBV) (18). *M.tb* purified protein derivative (PPD) was included as a positive control indicative of mycobacterial exposure. Initial crude analyses revealed evidence of differences in the levels of antibodies specific to PPD, DT, RSV, MV, KSHV ORF73, KSHV 8.1 and CMV antigens across the groups (Fig. 1). There were also differences in total IgG antibody concentrations (Supplementary Fig. 1). There was no evidence of differences in EBV and adenovirus-specific antibodies across the groups (Supplementary Fig. 1).

**Figure 1:**
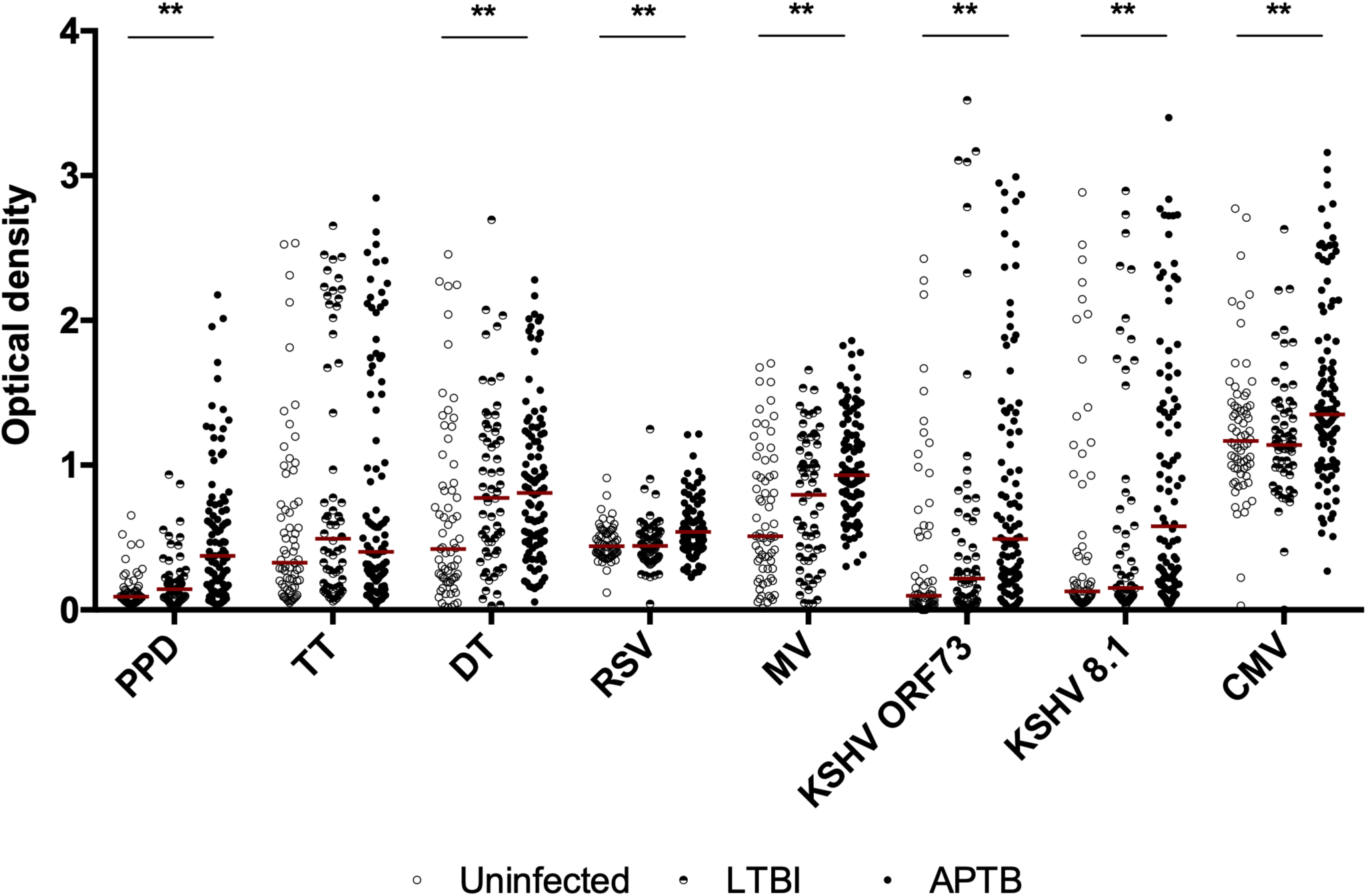
Variations in antibody responses to heterologous pathogens across *M.tb* infection state. The horizontal bars shown are median IgG antibody optical densities in each group. Antibody responses were compared across uninfected controls (n=68), individuals with LTBI (n=62) and APTB cases (n=107). The p values shown correspond to results from Kruskal–Wallis test (*p<0.05, **p<0.01). PPD: purified protein derivative, TT: tetanus toxoid, DT: diphtheria toxoid, RSV: respiratory syncytial virus, MV: measles virus, KSHV: Kaposi’s sarcoma herpesvirus, CMV: cytomegalovirus

In the final analyses, associations between *M.tb* infection status and antibody responses were determined by means of linear regression analysis using the uninfected controls as the baseline comparison group while adjusting for the effects of age, gender, HIV infection status and socioeconomic status. APTB was associated with higher levels of anti-PPD, anti-TT, anti-DT, anti-RSV, anti-MV and anti-KSHV 8.1 compared to the uninfected individuals but there was no association between *M.tb* infection status and anti-KSHV ORF73 or anti-CMV antibodies (Table 1). Total IgG levels were raised in the APTB cases compared to the uninfected controls (Supplementary Table 2). There were no associations between *M.tb* infection status and anti-EBV or anti-adenovirus antibodies, in agreement with findings from the crude analyses (Supplementary Table 2). LTBI was associated with higher anti-DT antibodies and weakly associated with higher anti-TT antibodies. LTBI was also marginally associated with higher anti-PPD antibodies. Taken together, these results show that *M.tb* infection, particularly APTB, is associated with higher levels of antibody responses to several heterologous pathogen recall antigens.

**Table 1:** Associations between *Mycobacterium tuberculosis* infection status and concentrations of IgG antibodies to heterologous pathogen antigens *.

| Antibody optical density | Adjusted GMR (95%CI)<sup>†</sup> | P value |
| --- | --- | --- |
| **Anti-PPD** |  |  |
| Uninfected | 1 |  |
| LTBI | 1.286 (0.995 - 1.663) | 0.055 |
| APTBI | **3.326 (2.446 - 4.524)** | **<0.001** |
| **Anti-TT** |  |  |
| Uninfected | 1 |  |
| LTBI | 1.448 (0.969 - 2.164) | 0.071 |
| APTBI | **1.608 (1.082 - 2.390)** | **0.019** |
| **Anti-DT** |  |  |
| Uninfected | 1 |  |
| LTBI | **1.443 (1.002 - 2.079)** | **0.049** |
| APTBI | **1.516 (1.123 - 2.047)** | **0.007** |
| **Anti-RSV** |  |  |
| Uninfected | 1 |  |
| LTBI | 0.922 (0.806 - 1.053) | 0.230 |
| APTBI | **1.178 (1.066 - 1.301)** | **0.001** |
| **Anti-MV** |  |  |
| Uninfected | 1 |  |
| LTBI | 1.074 (0.798 - 1.445) | 0.639 |
| APTBI | **1.599 (1.286 - 1.988)** | **<0.001** |
| **Anti-KSHV ORF73** |  |  |
| Uninfected | 1 |  |
| LTBI | 1.051 (0.926 - 1.192) | 0.441 |
| APTBI | 1.142 (0.994 - 1.312) | 0.062 |
| **Anti-KSHV K8.1** |  |  |
| Uninfected | 1 |  |
| LTBI | 1.112 (0.695 - 1.778) | 0.658 |
| APTBI | **1.969 (1.201 - 3.227)** | **0.007** |
| **Anti-CMV** |  |  |
| Uninfected | 1 |  |
| LTBI | 0.893 (0.615 - 1.298) | 0.553 |
| APTBI | 1.029 (0.782 - 1.354) | 0.837 |

GMR: geometric mean ratio, LTBI: latent tuberculosis infection, APTBI: active pulmonary tuberculosis, PPD: purified protein derivative, TT: tetanus toxoid, DT: diphtheria toxoid, RSV: respiratory syncytial virus, MV: measles virus, KSHV: Kaposi's sarcoma herpesvirus, CMV: cytomegalovirus
\* Linear regression analysis of antibody data from 67 uninfected controls, 62 individuals with LTBI and 89 APTBI cases.
<sup>†</sup> Adjusted for age, gender, socioeconomic status and HIV infection status.

### 2.2 BCG vaccination is not associated with increased antibody responses to unrelated pathogens

Next, we determined whether recent vaccination with the related species, *Mycobacterium bovis* BCG, was also associated with elevated antibody responses to unrelated recall pathogen antigens. Antibody responses were evaluated in a UK cohort of adolescents comprising a test group of 12 individuals who received BCG, and 13 age-matched BCG naïve controls, and the two compared at timepoints before and 3 weeks after vaccination of the test group. This cohort allowed us to compare BCG vaccinated individuals to a BCG naïve group, which we could not do in the Ugandan participants because BCG is routinely given to infants as part of the Uganda National Expanded Programme on Immunisation (19).

We observed no differences in antibody responses to any antigen, including PPD, between the BCG vaccinated individuals and naïve controls at the 3 week time point (Figure 2a). These results mirrored non-significant differences observed between the two groups before BCG vaccination (Figure 2b). These findings indicate that BCG vaccination may not be able to boost antibody responses to heterologous pathogens, or at least not within the 3 week time frame.

**Figure 2:**
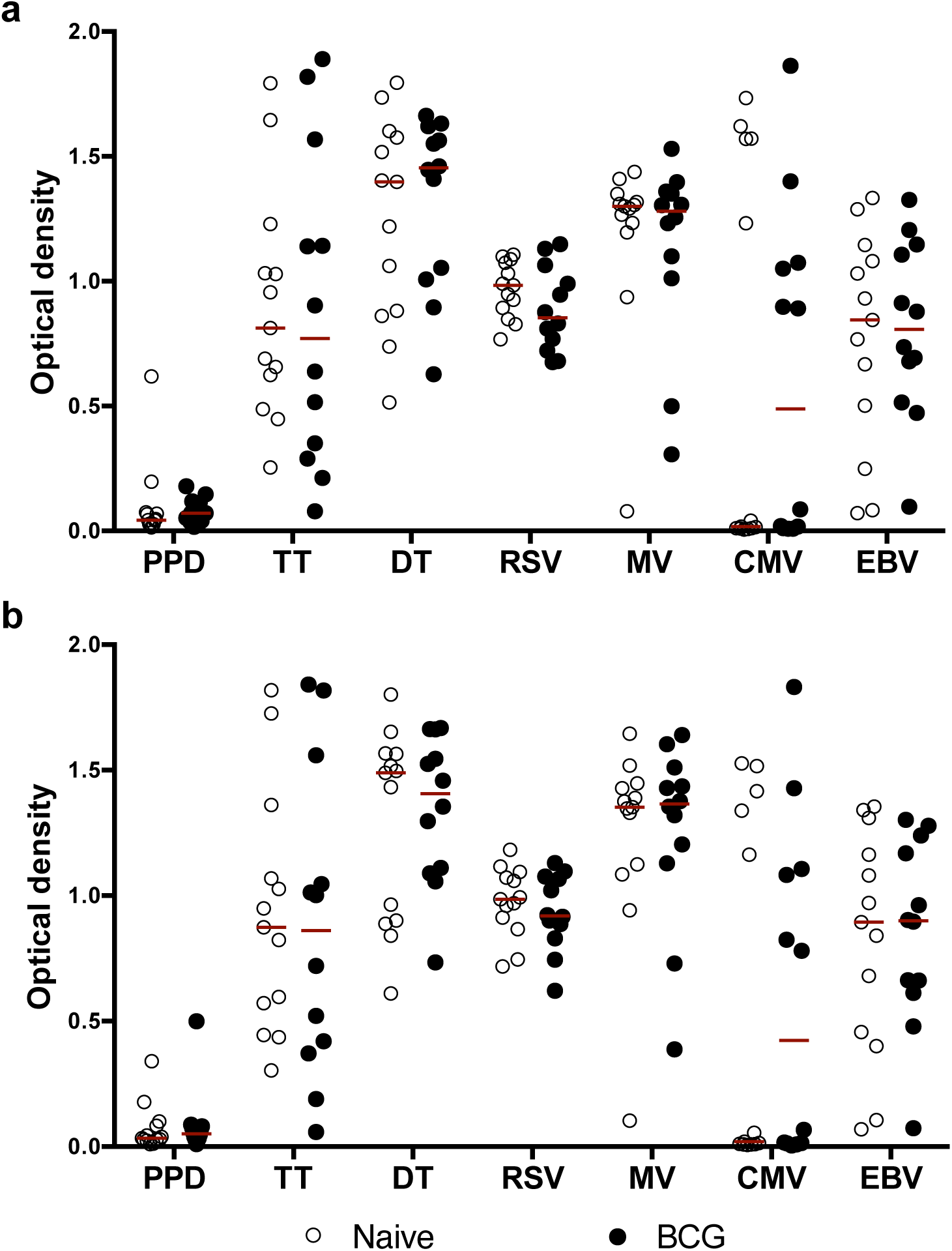
Antibody responses to heterologous pathogens in BCG vaccinated individuals and their age-matched BCG naïve controls. Panel a: antibody responses 3 weeks after BCG vaccination. Panel b: antibody responses before BCG vaccination. The horizontal bars shown are median IgG antibody optical density in each group. The p values shown correspond to results from Wilcoxon rank sum test (*p<0.05, **p<0.01) from comparing antibody responses in BCG vaccinated (n=12) and BCG naïve controls (n=13). PPD: purified protein derivative, TT: tetanus toxoid, DT: diphtheria toxoid, RSV: respiratory syncytial virus, MV: measles virus, CMV: cytomegalovirus, EBV: Epstein Barr virus.

### 2.3 Avidity of tetanus toxoid and measles virus-specific antibodies is increased in APTB

We determined whether the observed increases in antibody responses to unrelated recall antigens in *M.tb* infected individuals was accompanied by changes in their avidity or overall binding strength. An increase in avidity could point to MBCs being the source of heterologous pathogen antigen-specific antibodies because these cells would have already undergone affinity maturation. To this end, the avidity of antibodies against tetanus toxoid (TT) and measles virus haemagglutinin (MVHA) antigens were investigated using SPR. Due to limitations in sample volumes, only 88 of the 237 KTB study participants’ samples were tested.

Initial crude comparisons showed marginal evidence of a difference in dissociation rate (the measure of avidity) in TT-specific antibodies across the three groups (Figure 3). For the final analysis, linear regression was used to evaluate associations between *M.tb* infection state and antibody avidity using the uninfected controls as the baseline comparison group while adjusting for the effects of age, gender, HIV infection status and socioeconomic status (Table 2). APTB and LTBI were associated with the presence of TT-specific antibodies of a slower dissociation rate, a sign of higher avidity. Having APTB was also associated with MVHA specific antibodies of a slower dissociation rate. These results lend credence to the hypothesis that the increased antibody responses against heterologous pathogen recall antigens observed in *M.tb* infection arise from antigen-specific MBCs.

**Table 2:** Association between *M.tb* infection state and SPR derived TT and MVHA specific antibody dissociation rates ^‡^.

| Dissociation rate E-06 [kd(s-1)] | Adjusted GMR (95%CI) § | Pvalue |
| --- | --- | --- |
| **Anti-TT antibodies** |  |  |
| Uninfected | 1 |  |
| LTBI | **0.728 (0.548 - 0.967)** | **0.029** |
| APTB | **0.725 (0.582 - 0.902)** | **0.004** |
| **Anti-MVHA antibodies** |  |  |
| Uninfected | 1 |  |
| LTBI | 0.971 (0.915 - 1.030) | 0.331 |
| APTB | **0.927 (0.861 - 0.998)** | **0.044** |

GMR: geometric mean ratio, LTBI: latent tuberculosis infection, APTB: active pulmonary tuberculosis
‡ 23 uninfected controls, 25 individuals with LTBI and 34 APTB cases
§ Adjusted for age, gender, socioeconomic status and HIV infection status

**Figure 3.**
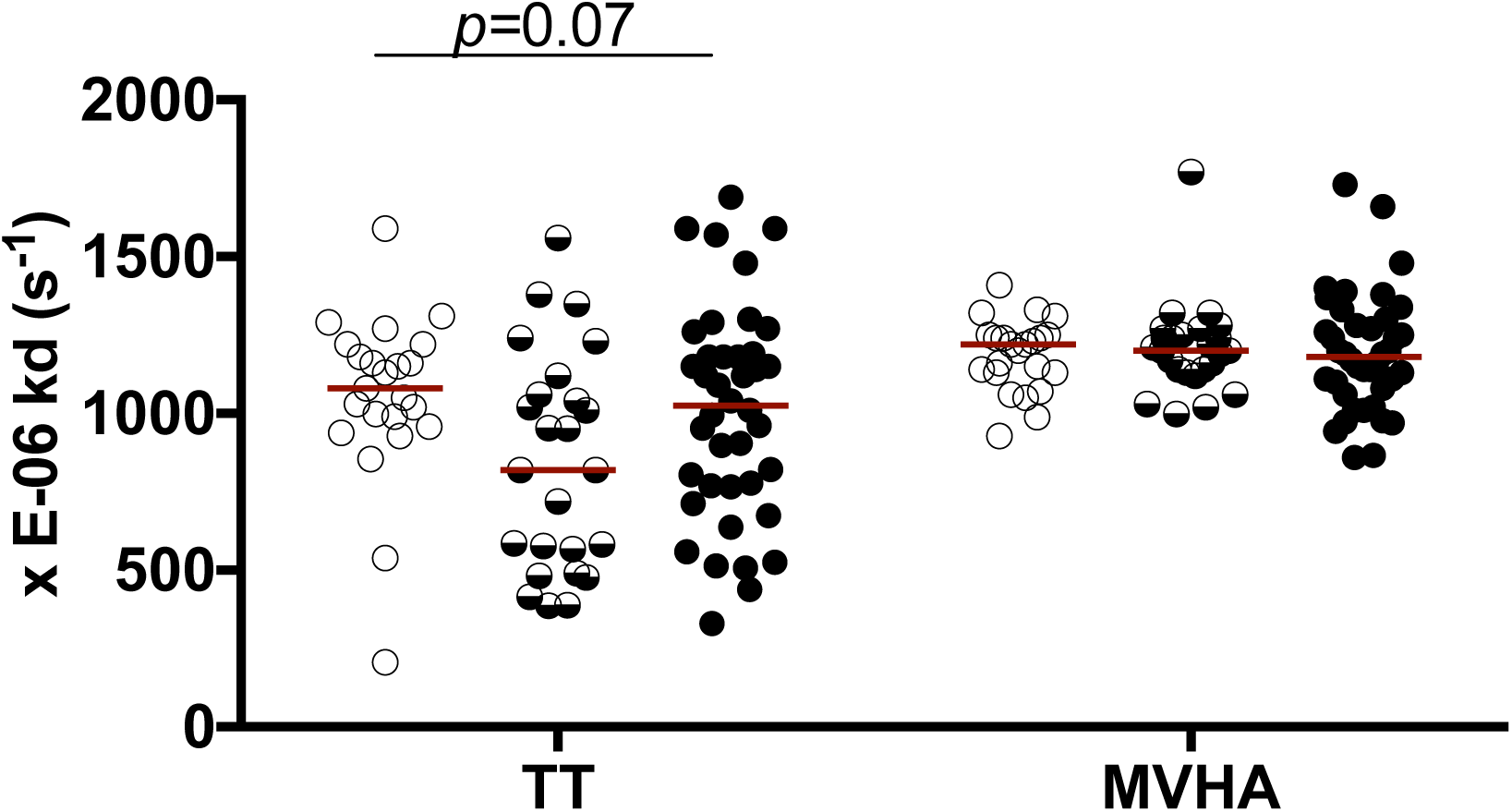
Variation in tetanus toxoid and measles virus-specific antibody avidity across M.tb infection state. The Kruskal Wallis test was used to compare antibody dissociation rates across uninfected controls (n=23), individuals with LTBI (n=25) and APTB cases (n=40). TT: tetanus toxoid, MVHA: measles virus haemagglutinin

### 2.5 Tetanus toxoid-specific plasmablasts are higher than tetanus toxoid-specific memory B cells in APTB

In order to explore the possibility that *M.tb* infection may be driving differentiation of MBCs specific to unrelated recall antigens into plasmablasts in our study participants, we examined the frequency of TT-specific MBCs and plasmablasts in 48 APTB cases and 115 healthy donors using B cell ELISPOT assays.

We had limited PBMC numbers from the APTB cases and so MBC frequencies were ascertained in 18 of the APTB cases while plasmablast frequencies were evaluated in the rest (n=30). The ratio of the median frequencies of TT-specific plasmablasts to MBC in the APTB cases was 7:1 (8.75:1.25; Figure 4). The high abundance of TT-specific plasmablasts relative to MBCs in the APTB cases provides evidence of polyclonal activation of antigen-specific MBCs in APTB. This process may be driving their differentiation into plasmablasts and could explain the increased levels of antibodies against heterologous pathogen recall antigens observed in active TB disease.

**Figure 4.**
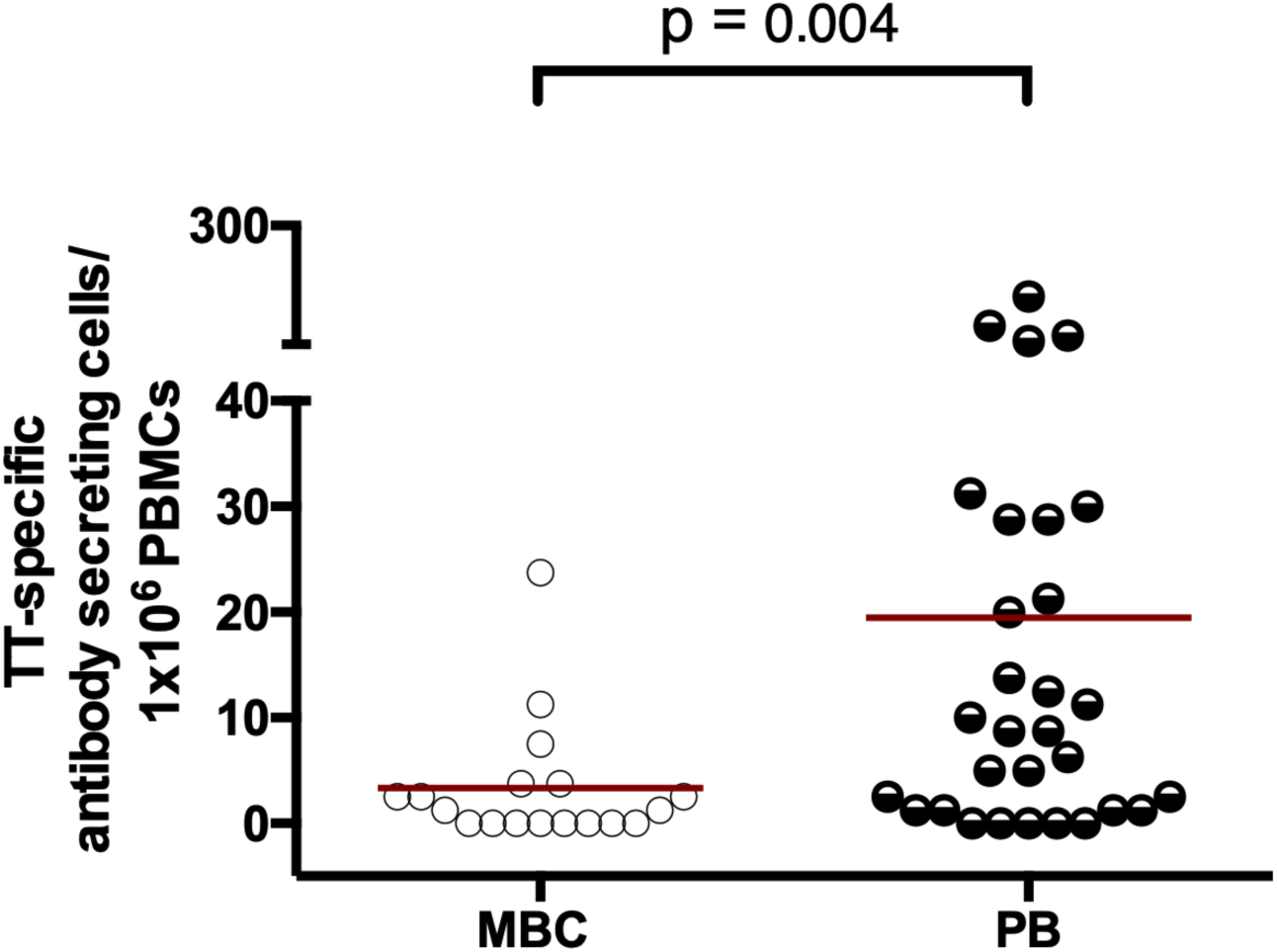
Tetanus toxoid-specific plasmablasts are higher than tetanus toxoid-specific memory B cells in APTB. MBC: memory B cells, PB: plasmablasts. MBC responses were evaluated in 18 APTB cases while PB responses were evaluated in 30 APTB cases. The p values are from Wilcoxon-rank sum tests. TT: tetanus toxoid

Efforts to determine TT-specific plasmablast to MBC ratio in the healthy donors for comparison were not possible because TT-specific plasmablasts were not readily detected during preliminary analyses. However, high frequencies of TT-specific MBC were detected in these individuals at levels above those seen in the APTB cases (Supplementary Fig. 2). Although TT-specific MBCs have been shown to remain in circulation years after vaccination, TT-specific plasmablasts are short-lived, disappearing within 2 weeks after antigen exposure (20). This may explain the lack of detectable TT-specific plasmablasts in healthy donors.

## 3.0 Discussion

This study highlights for the first time a role for natural *M.tb* infection in the boosting of antibody immunity to heterologous pathogens that individuals may have been previously exposed to, through infection or immunisation, and shows that vaccination with the related bacterium, *Mycobacterium bovis* BCG does not have a similar effect. The increase in the levels and avidity of these antibodies in *M.tb* infected individuals means that MBCs are their likely source. Additionally, the abundance of TT-specific plasmablasts relative to TT-specific MBCs in active TB disease provides evidence to support the hypothesis that *M.tb* infection drives polyclonal activation of MBCs generated against unrelated recall antigens. If this property can be exploited, it may lead to the production of immunological therapies than can be used to boost serological memory to a wide variety of infections, particularly in people with waning immunity such as the elderly.

The fact that the increase in antibody responses to heterologous pathogens was most evident in APTB disease may mean that this non-specific effect is most pronounced in a milieu of actively replicating tubercle bacilli. Indeed, previous research has shown that active TB disease is associated with increases in cytomegalovirus-specific antibodies (21,22), however, these studies attributed the rise in antibodies to concurrent infection with CMV rather than as a marker of serological memory. A study by de Paus and colleagues (2013) observed higher levels of influenza-specific antibodies in individuals with TB and recognized that in addition to concurrent infection there was a possibility that these may have arisen as a result of mycobacterial driven non-specific increases in antibodies. The marginal increases in TT-specific antibody levels in LTBI and the statistically significant increase in SPR determined TT-specific antibody avidity in LTBI indicate that LTBI, like APTB, may boost antibody responses to heterologous pathogens, albeit to a lesser extent. It is now recognised that LTBI is not a fully quiescent state of infection but may at times be associated with subclinical disease arising from reactivation of *M.tb* (24,25). Mycobacterial growth in this state may drive the observed non-specific increases in antibody responses.

The observation that BCG vaccination did not have a similar effect may mean that proteins unique to *M.tb* may mediate the increase in antibody responses to these heterologous pathogens observed in APTB. Alternatively, this may point to the effect of BCG mycobacterial load at the site of inoculation, which may fall short of the *M.tb* bacillary load observed in the lungs during active TB disease. The lack of significant stimulation of anti-mycobacterial antibodies in the BCG vaccinated recipients, in contrast to the more than 3-fold rise in the same in the APTB cases (Table 1) supports this reasoning. The lack of antibody response may also simply be due to the timing after vaccination. A later time point may have allowed the accrual of BCG-specific antibodies; we were not able to assess this in our study.

Antibody responses to recall antigens from heterologous pathogens were increased in both level and avidity in *M.tb* infection. This gave further evidence that the source of these antibodies are MBCs because these cells would have already undergone affinity maturation, and as a consequence produce antibodies of high avidity. A study by Zimmerman and colleagues (2016) argues that the primary source of anti-mycobacterial antibodies in TB disease are MBCs, as demonstrated by the high level of somatic hypermutations in antibody-secreting cells isolated in this disease state. Tubercle bacilli may drive the activation of MBCs specific to a wide variety of pathogen antigens including those specific for *M.tb*. A similar phenomenon has been demonstrated for other infections such as malaria (27), hepatitis C (28) and hantavirus pulmonary disease (29). Bernasconi and colleagues (2006) have previously shown that MBCs are easily activated by polyclonal stimuli such as bacterial DNA and the cytokines evoked following vaccination. It is, therefore, possible that mycobacterial products and cytokine milieu present during human *M.tb* infection may activate MBCs leading to their differentiation to antibody-secreting cells. Our B cell ELISPOT data illustrated an abundance of TT-specific plasmablasts relative to TT-specific MBCs in APTB. Frequencies of TT-specific MBCs were lower in APTB cases in comparison to healthy donors while TT-specific plasmablasts were not readily detected in the latter. These observations further support the hypothesis that *M.tb* drives MBC polyclonal activation. It is not clear how the *M.tb* may benefit from increasing recall antibody responses to other pathogens. However, we speculate that this process may aid in the survival of *M.tb* within the host by limiting host death as a result of infection by unrelated pathogens.

It is unclear as to why APTB was only significantly associated with antibody responses to some pathogen recall antigens and not others. This may relate to varied rates of exposure to the different infections over time, with multiple recent exposures masking any “benefit” gleaned from mycobacterial driven MBC activation. Certain pathogens such as EBV has been shown to establish latency in memory B cells (31). This virus has also been shown to increase proliferation and activation of B cells and add to the effect of known TLR agonists such as CpG (32). This would support the idea of increased EBV antibody responses in *M.tb* infected individuals, however, this was not observed in our study.

This study has a few limitations. We assumed that the antibody responses detected against the chosen heterologous pathogen antigens in our study participants arose as a result of past exposure via infection or vaccination. However, recent or current exposure to these multiple pathogens, although unlikely, was possible.

On the whole, these data show a hitherto unknown role for *M.tb* in the maintenance of serological memory generated against past infections and vaccinations. It has been hypothesised that exposure to infections in later life could help maintain antibody responses to a broad set of pathogens in the absence of their specific antigens. This process is proposed to be due to polyclonal stimulation of antigen MBCs (11,30) and the data presented here would support this. However, further experimentation is required to confirm the mechanism involved and identify the specific *M.tb* antigens mediating this process. This research may aid in the development of therapeutics to boost antibody immunity to heterologous pathogens. It is envisaged that this treatment could be useful in ageing populations in whom antibody responses to specific infections are known to wane. The advantage of this approach is that instead of boosting these individuals with multiple vaccine antigens, a single treatment could be provided to increase serological immunity to several pathogens all at once.

## 4.0 Materials and Methods

### 4.1 Study populations and design

A cross-sectional study design was used to determine whether there were associations between *M.tb* infection status and antibody responses to heterologous pathogen recall antigens in study participants from a TB household contact study based in Uganda investigating the effect of coinfections in humans on their susceptibility to infection with *M.tb* (33). Individuals with APTB and their household contacts (HHCs) were recruited from the suburbs of Kampala. The APTB cases were all adults over 18 years of age and had either only recently began anti-TB treatment or received treatment for less than 4 weeks. APTB was ascertained using acid fast bacilli sputum microscopy. Individuals with LTBI were identified from among healthy HHCs using the tuberculin skin test and QuantiFERON-TB Gold in-tube (QFT-GIT) test. The HHCs that tested positive on both these tests were classified as having LTBI whereas those that tested negative on both tests were classified as uninfected controls. A total of 68 uninfected controls, 62 individuals with LTBI and 107 APTB cases were included in our study. QuantiFERON-TB Gold in-Tube (QFT-GIT) nil supernatant samples from these individuals were used for the antibody analyses. The choice of QFT-GIT supernatants was because serum samples were not available for all the individuals studied. We have previously shown that there are no differences between antibody concentrations obtained from assaying serum and those from assaying QFT-GIT supernatants (34).

Stored plasma samples from adolescents participating in a UK based prospective study on BCG responses were used to determine whether BCG vaccination was associated with non-specific increases in antibody responses to heterologous pathogen recall antigens. These individuals were part of a UK government programme on BCG vaccination in schools in the year 2005 and were invited to take part in a research study that sought to identify immune responses involved in BCG mediated protection against TB. The adolescents studied included 12 subjects who were given BCG and then followed up after three weeks as well as 13 age-matched controls who did not receive BCG and were followed up for the same period of time. All the subjects had no previous BCG scar (a sign of earlier BCG vaccination) and had a negative TST prior to immunisation (13). Samples obtained from these individuals before BCG vaccination and at the 3 week timepoint were analysed. This cohort was chosen because it gave us the unique opportunity to compare BCG vaccinated and unvaccinated controls, something which could not be easily done in Ugandan cohorts considering BCG is part of the immunisation schedule in the country (19).

In order to investigate MBC polyclonal activation as a possible cause of the elevated antibody responses to heterologous pathogens in people with TB, cryopreserved PBMCs from 48 APTB cases from the TB household contact study and fresh PBMCs from 115 healthy donor controls were analysed to determine the effect of an active TB infection on the frequencies of tetanus toxoid-specific plasmablasts and MBCs. The healthy donors were recruited from a voluntary HIV counselling and testing clinic at the Uganda Virus Research Institute. They were all HIV negative, aged 18-55 years and found to be healthy by a clinician.

These studies were all exploratory and the sample size was determined by the availability of sufficient samples and reagents for laboratory analyses.

### 4.2 Ethics Statement

Informed written consent was obtained from all the adult study participants. The parents of the adolescent participants provided written informed consent and verbal consent was obtained from the participants themselves. This research was approved by Research & Ethics Committees at the Makerere University School of Biomedical Sciences and School of Medicine, the Uganda Virus Research Institute, the London School of Hygiene & Tropical Medicine and Uganda National Council for Science and Technology.

### 4.3 Laboratory assays

#### 4.3.1 In-house antigen-specific IgG ELISA

Immulon 4 HBX microtiter plates (Thermo Scientific, USA) were coated with 50 µl/well of 5 µg/ml diphtheria toxoid, 5 µg/ml tetanus toxoid (both from National Institute for Biological Standards and Control), purified protein derivative (PPD) of *M.tb* (Serum Statens Institute, Denmark), respiratory syncytial virus antigen, measles grade 2 antigen, adenovirus grade 2 antigen, 1.25 µg/ml Epstein Barr viral capsid (all from Microbix, Biosystems, Canada) or 0.1% (w/v) skimmed milk (control for nonspecific binding) in carbonate-bicarbonate buffer (pH 9.6) at 4 °C overnight. Following overnight incubation, the plates were washed four times with 1x PBS (pH 7.4) containing 0.05% Tween 20 (PBS-T). The plates were then blocked with 150 µl/well 1% (w/v) skimmed milk in PBS-T for 2 h at room temperature. A dilution of 1 in 100 of each sample in 0.1% skimmed milk PBS-T (assay buffer) was made and 50 µl was then added to the antigen-coated and control wells. After an incubation of 2 h at room temperature, the wells were washed as before and incubated with 50 µl/well polyclonal rabbit anti-human IgG conjugated with horseradish peroxidase (Dako, Denmark) at 0.5 µg/ml for another hour at room temperature. Plates were then washed four times and enzyme activity detected by incubation with 100 µl/well o-phenylenediamine (Sigma) containing hydrogen peroxide for 15 min at room temperature in the dark. The reaction was stopped by addition of 25 µl/well 2 M sulphuric acid and thereafter the optical density (OD) measured at a test wavelength of 490 nm and a reference wavelength of 630 nm in an ELISA plate reader (Biotek). The ODs from the control wells were subtracted from the test antigen wells to eliminate background antibody levels.

#### 4.3.2 KSHV antibody ELISA

Immulon 4 HBX microtiter plates pre-coated with K8.1 or ORF73 were kindly provided by the Viral Oncology Section, AIDS and Cancer Virus Program, SAIC-Frederick, Inc., NCI-Frederick, Frederick, MD 21702. In the coating procedure, 100 µl/well of 2 µg/ml K8.1 or ORF73 diluted in 0.05 M carbonate/bicarbonate buffer, pH 10 or 1x PBS respectively was incubated in plate wells overnight at 4 °C. The plates were washed three times with PBS-T and then blocked with 280 µl/well of assay buffer [2.5% (w/v) BSA (Sigma), 2.5% (v/v) normal donor goat serum (Equitech-Bio), 0.005% (v/v) Tween 20, 0.005% (v/v) Triton X-100 in 1x PBS] for 3 h at 37 °C and thereafter stored at 80 °C until use.

The plates were thawed at 37 °C in preparation for the assay and washed three times with PBST. A 1 in 100 dilution of the sample or controls was prepared and then 100 µl was added per well in duplicate. Each plate had 3 positive and 2 negative control samples for quality control. After an incubation of 90 min at 37 °C, the wells were washed five times and incubated with 100 µl/well goat anti-human IgG conjugated with alkaline phosphatase (Roche Diagnostics) diluted 1 in 5,000 for 30 min at 37 °C. The plates were washed five times and enzyme activity detected by incubation with 100 µl/well 1-step p-nitrophenyl phosphate substrate solution (Thermo Scientific Pierce, USA) for 30 min at room temperature in the dark. This reaction was stopped by the addition of 50 µl/well of 3 N NaOH and then the plates were read at a wavelength of 405 nm.

#### 4.3.3 CMV antibody ELISA

Platelia™ CMV IgG ELISA was used to measure CMV-specific antibodies as per the manufacturer’s instructions (3). Briefly, diluted samples, standards of known anti-CMV antibody concentration and controls were added to microtitre plates pre-coated with CMV antigens. The plates were incubated with the samples at 37°C for 45 min and thereafter washed with buffer. A conjugate composed of horseradish peroxidase enzyme bound onto anti-human IgG monoclonal antibody was then added to each well and incubated for 45 min at 37°C. The plates were washed and an enzyme substrate was added to each well followed by a 15-min incubation at room temperature. After this, a stop solution was added and the OD read at 450 nm wavelength.

#### 4.3.4 Total IgG antibody ELISA

Human IgG total Ready-SET-Go (Affymetrix ebioscience) was used to measure total IgG following the manufacturer’s instructions (35). In the procedure, Immulon 4 HBX microtiter plates were coated with 100 µl/well purified anti-human IgG monoclonal antibody overnight at 4 °C. The plates were then washed twice with wash buffer and incubated with 250 µl/well blocking buffer for 2 h at room temperature. The plates were washed twice and eight, recombinant human IgG controls were prepared and the samples were diluted 1 in 100,000. A volume of 100 µl/well of the prepared standards and samples was added to the plates and then incubated for 2 h at room temperature. In the next step, the plates were washed four times and incubated with 100 µl/well HRP-conjugated anti-human IgG monoclonal detection antibody for 1 h at room temperature. The plates were washed four times and incubated with 100 µl/well tetramethylbenzidine (TMB) substrate solution for 15 min at room temperature in the dark. After this duration, the reaction was stopped by addition of 100 µl/well stop solution and the plates read at 450 nm wavelength.

#### 4.3.5 Surface plasmon resonance

The avidity of antibodies directed against tetanus toxoid and measles virus haemagglutinin (MVHA) antigens was evaluated. These antigens were immobilised onto CM5 sensor chips (Biacore, GE Healthcare, Amersham) by amine coupling to a level of 2000 RU.

Samples were diluted 1 in 3 in HBSPE and run through Bio-Gel® P-30 (Bio-Rad, UK) polyacrylamide gel spin columns to minimise non-specific binding. They were further diluted 1 in 8 in HBSPE running buffer containing 1% (w/v) carboxymethyl-dextran sodium salt (Sigma) and analysed in the Biacore 3000 instrument at a temperature of 25°C. A 90 µl volume of sample was injected over the chip surface at a rate of 15 µl/min followed by a dissociation time of 8 min. Prior to analysis of the next sample, the chips were regenerated with 50 mM HEPES containing 3 M MgCl_2_ and 25% (v/v) ethylene glycol, followed by 20 mM glycine pH 1.5 and re-equilibration in HBSPE.

BIAevaluation software version 4.1.1 was used for data analysis and control flow-cell traces with immobilised alpha-1 antitrypsin background were subtracted from test flow cell data. A Langmuir 1:1 dissociation model was used to determine the dissociation rate between 10 seconds and 300 seconds post sample injection.

#### 4.3.6 B cell ELISPOT

We used in-house *in vitro* and *ex-vivo* B cell ELISPOTS to determine the frequency of TT-specific plasmablasts and MBCs respectively in APTB cases and healthy donors following methods described by Sebina *et al.* (36,37). Cryopreserved PBMCs were thawed, resuspended in RPMI (Gibco by Life Technologies) containing 10% (v/v) fetal bovine serum, 2 mM L-glutamine (both from Sigma-Aldrich, UK) and 1x penicillin-streptomycin (Invitrogen). The cells were then rested for 4-6 hours at 37°C and 5% CO_2_.

In the *in vitro* ELISPOT, PBMCs at a cell density of 1×10^6^ cells/ml were stimulated with a mixture of 25 ng/ml human IL-10 (R&D Systems, UK), 0.5 µg/ml of pokeweed mitogen (PWM), 6 µg/ml of CpG oligodeoxynucleotide, 1.2 mg/ml *Staphylococcus aureus* Cowan, 50 µM ß-mercaptoethanol at 37°C and 5% CO_2_ for six days. Approximately 4×10^5^ of the cultured cells were then transferred to each well of 96-well filter (Merck Milipore) pre-coated with 2 µg/ml TT in 1x PBS or only 1xPBS. The plates were sealed and incubated for 6 hours at 37°C and 5% CO_2_ after which biotin-SP conjugated affinipure fragment donkey antihuman IgG (Jackson ImmunoResearch) was added to the plate wells to aid in the detection of anti-TT specific antibody secreting cells. After overnight incubation at 4°C, streptavidin-AKP (BD biosciences) was added and the plates incubated for an hour at room temperature followed by AP-conjugate substrate (Bio-rad, USA) to develop the spots. After 10 minutes the plate wells were rinsed with water and left to dry before they were read using an AID ELISPOT reader (AID Diagnostika, Germany). The number of MBCs was then expressed as the total number of spots per million cells.

The *ex vivo* ELISPOT involved the direct transfer of the rested PBMCs to the 96-well filter plates at a cell density of 4×10^5^ cells/well. All the subsequent steps were similar to those described for the *in vitro* ELISPOT.

### 4.4 Statistical Analysis

Results were analysed using Stata release statistical package and GraphPad Prism software. The initial analyses were made using the Kruskal-Wallis test to compare antibody responses across the groups of uninfected controls, individuals with LTBI and active TB cases. Linear regression analysis with bootstrap confidence intervals was then be used to determine associations between *M.tb* infection status and antibody responses (12,17). Adjusting was done for the effects of potential confounders such as HIV infection, age, socioeconomic status (SES) and gender. The Wilcoxon rank sum test was used to compare antibody responses between BCG vaccinated individuals and BCG naïve controls as well as frequencies of MBCs and PBs in APTB cases.

## Acknowledgements

This research was supported by Wellcome Trust Uganda PhD Fellowships in Infection and Immunity held by IAB and SGK, funded by Wellcome Trust Strategic Award Grant no. 084344 and a Wellcome Trust Masters Training Fellowship (Grant no. 092779) held by IS. SGK also received support from a Commonwealth Scholarship Commission Split Site PhD Fellowship and through the DELTAS Africa Initiative (Grant no. 107743). The DELTAS Africa Initiative is an independent funding scheme of the African Academy of Sciences (AAS), Alliance for Accelerating Excellence in Science in Africa (AESA), and supported by the New Partnership for Africa’s Development Planning and Coordinating Agency (NEPAD Agency) with funding from the Wellcome Trust (Grant no. 107743) and the UK Government. AME and SC received funding from Medical Research Council UK, grant number MR/K019708/1 and AME also received funding from the Wellcome Trust, grant number 095778. The views expressed in this publication are those of the author(s) and not necessarily those of AAS, NEPAD Agency, Wellcome Trust or the UK government.

## Supporting information

**Supplementary figure 1: Antibody responses to Epstein Barr virus and adenovirus and total IgG levels in individuals of varied *M.tb* infection status.** The horizontal bars shown are median IgG antibody optical densities in each group. Antibody responses against Epstein Barr virus (EBV) and adenovirus (panel a) at 1/100 sample dilution and total IgG levels (panel b) at a 1/100,000 sample dilution were compared across uninfected controls (n=68), individuals with LTBI (n=62) and APTB cases (n=107). The p values shown correspond to results from Kruskal–Wallis test (*p<0.05, **p<0.01). EBV: Epstein Barr virus

**Supplementary figure 2: Lower TT-specific Memory B cell (MBC) frequencies in APTB cases compared healthy donor controls.** MBC responses were evaluated in 115 healthy donors and 18 APTB cases. The p values are from Wilcoxon-rank sum tests. TT: tetanus toxoid

