## Supplementary Figure 1 and 2 for "*Mycobacterium tuberculosis* infection boosts B cell responses to unrelated pathogens"

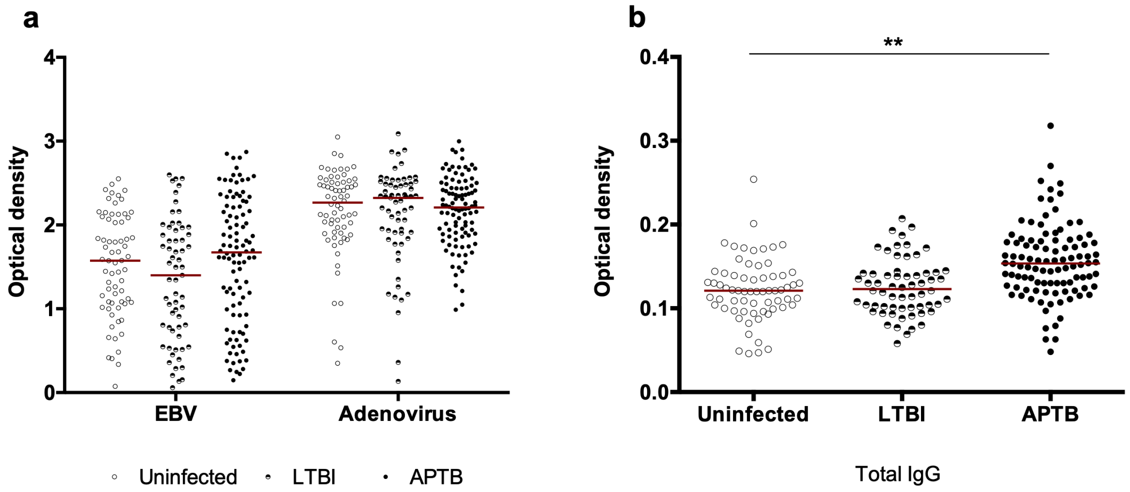


**Supplementary figure 1: Antibody responses to Epstein Barr virus and adenovirus and total IgG levels in individuals of varied *M.tb* infection status.** The horizontal bars shown are median IgG antibody optical densities in each group. Antibody responses against Epstein Barr virus (EBV) and adenovirus (panel a) at 1/100 sample dilution and total IgG levels (panel b) at a 1/100,000 sample dilution were compared across uninfected controls (n=68), individuals with LTBI (n=62) and APTB cases (n=107). The p values shown correspond to results from Kruskal–Wallis test (*p<0.05, **p<0.01). EBV: Epstein Barr virus

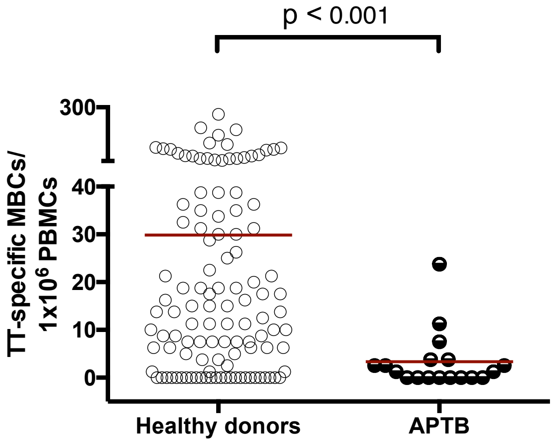


**Supplementary figure 2: Lower TT-specific Memory B cell (MBC) frequencies in APTB cases compared healthy donor controls.** MBC responses were evaluated in 115 healthy donors and 18 APTB cases. The p values are from Wilcoxon-rank sum tests. TT: tetanus toxoid
