## Supplementary Table 1 and 2 for "*Mycobacterium tuberculosis* infection boosts B cell responses to unrelated pathogens"

**Supplementary table 1: Study participant characteristics**

| Characteristic | Uninfected (n=68) | LTBI (n=62) | APTB (n=107) | *P* value |
|---|---|---|---|---|
| **Mean age & range (years)** | 16 (1, 66) | 24 (1, 66) | 30 (18, 53) | <0.0001 |
| **Females** | 39 (57.4%) | 41 (65.0%) | 44 (41.1%) | 0.006 |
| **HIV positive** | 6 (8.8%) | 5 (7.9%) | 42 (39.3%) | <0.0001 |
| **Low SES [b]** | 34 (50.8%) | 32 (51%) | 58 (61.0%) | 0.309 |

LTBI = latent tuberculosis infection, SES = socioeconomic status, and APTB = active pulmonary tuberculosis

*P* values are from chi-square tests of associations

[b] Individuals were either of low or medium SES

**Supplementary table 2: Associations between *Mycobacterium tuberculosis* infection status and IgG antibody responses against EBV and adenovirus antigens and total IgG** [*]

| Antibody optical density | Adjusted GMR (95%CI) [†] | *P* value |
|---|---|---|
| **Anti-EBV** | | |
| **Uninfected** | 1 | |
| **LTBI** | 0.802 (0.615 - 1.045) | 0.102 |
| **APTB** | 0.940 (0.727 - 1.216) | 0.638 |
| **Anti-adenovirus** | | |
| **Uninfected** | 1 | |
| **LTBI** | 0.928 (0.778 - 1.107) | 0.408 |
| **APTB** | 0.985 (0.892 - 1.089) | 0.769 |
| **Total IgG** | | |
| **Uninfected** | 1 | |
| **LTBI** | 1.027 (0.926 - 1.140) | 0.610 |
| **APTB** | **1.166 (1.038 - 1.309)** | **0.010** |

GMR: geometric mean ratio, LTBI: latent tuberculosis infection, APTB: active pulmonary tuberculosis, PPD: purified protein derivative, TT: tetanus toxoid, DT: diphtheria toxoid, RSV: respiratory syncytial virus, MV: measles virus, KSHV: Kaposi's sarcoma herpesvirus, CMV: cytomegalovirus, EBV: Epstein Barr virus.

[*] Linear regression analysis of antibody data from 67 uninfected controls, 62 individuals with LTBI and 89 APTB cases.

[†] Adjusted for age, gender, socioeconomic status and HIV infection status.
